# Social isolation impairs the prefrontal-nucleus accumbens circuit subserving social recognition in mice

**DOI:** 10.1101/2020.10.08.332320

**Authors:** Gaeun Park, Changhyeon Ryu, Soobin Kim, Yong-Seok Lee, Sang Jeong Kim

## Abstract

The medial prefrontal cortex (mPFC) plays important roles in social behaviors, but it is not clear how early social experiences affect the mPFC and its subcortical circuit. We report that mice singly housed for 8 weeks immediately after weaning (SH mice) show a deficit in social recognition, even after 4 weeks of re-socialization. In SH mice, prefrontal infralimbic (IL) neurons projecting to the shell region of nucleus accumbens (NAcSh) showed decreased excitability compared to normally group housed (GH) mice. Furthermore, NAcSh-projecting IL neurons were activated when the mice encountered a familiar conspecific, which was not shown in SH mice. Chemogenetic inhibition of NAcSh-projecting IL neurons in normal mice selectively impaired social recognition without affecting social interaction, whereas activation of these neurons reversed social recognition deficit in SH mice. Therefore, mPFC IL-NAcSh projection is a novel brain circuit affected by early social experience; its activation is required for the social recognition.

## Introduction

Social behaviors include a wide range of behaviors including social preference and social recognition^1, 2, 3^. Social recognition is the behavioral characteristics of distinguishing a novel conspecific from a familiar one, which is evolutionarily well-conserved due to its importance for survival^4, 5, 6^. Social recognition is impaired in several neurodevelopmental disorders such as autism spectrum disorder (ASD) and schizophrenia^7, 8, 9^. Although several brain regions, including the hippocampus, have been suggested as a hub for social recognition, the neural circuit mechanism subserving social recognition remains largely unknown^10, 11^.

Early social experience is known as one of the key environmental factors for postnatal neurodevelopments^12, 13, 14^. In particular, social deprivation during the critical developmental period has the long lasting effect on social function in adults^15, 16, 17, 18^. Juvenile social isolation (jSI) is one of animal models widely used to study the effects of early social experiences on postnatal developments and social functions in adulthood. In mice, jSI was shown to result in social behavior deficits in adulthood; however, previous studies using the jSI model have mostly focused on its effect on social preference^19, 20^, whereas the aspects of social behaviors that are more sensitive to jSI have not been fully investigated. In addition, whether the selective deficits, if there are any, can be reversed by resocialization remains controversial^21, 22, 23, 24^.

The medial prefrontal cortex (mPFC) is critically involved in social behaviors in both humans and rodents. Imaging studies in humans and mice showed that the mPFC is activated in response to social behaviors^25, 26, 27, 28^. Optogenetic manipulation of excitation/inhibition balance in the mPFC impairs social interaction in mice^29^. Importantly, the mPFC is one of the brain regions affected by jSI^18, 19, 30, 31, 32^. Early social isolation affects brain structures and physiology, as well as social behaviors^30, 33^. Recent studies have begun to reveal the mPFC-subcortical circuits affected by jSI. For example, Yamamuro and colleagues showed that mPFC neurons projecting to the posterior paraventricular thalamus (pPVT) are activated by social contacts and its activities are reduced in jSI mice^19^. However, it is not known whether the mPFC neurons projecting to the other subcortical regions implicated in social behavior, such as the nucleus accumbens (NAc), are also affected by jSI.

In this study, we found that social recognition is severely impaired by jSI, which was not recovered by a prolonged resocialization. We revealed that mPFC neurons projecting to the NAc shell (NAcSh) encode neural information for social familiarity, and the pharmacogenetic manipulation of this neuronal population selectively affects social recognition.

## Results

### Single housed mice showed a social recognition deficit after resocialization

A previous study showed that resocialization for 4 weeks was sufficient to restore the social interaction impairment induced by 8 weeks of social isolation^30^. We examined whether other social phenotypes are also affected by this isolation-regrouping protocol (Fig. 1a). We used the three-chamber test to test social preference and social recognition^4^ (Fig. 1b). In the social preference test, SH mice spent significantly more time exploring a conspecific than an object, showing that the social preference is, at least partially, restored in SH mice after resocialization (Fig. 1c). However, SH mice did not spend significantly more time interacting with novel conspecifics compared to the familiar conspecifics while GH mice spent significantly more time interacting with novel conspecifics, suggesting a severe deficit in social recognition in SH mice (Fig. 1d). Importantly, SH mice showed a comparable performance to GH mice in the novel object recognition test, demonstrating that SH mice do not have a general recognition memory deficit (Fig. 1e). Furthermore, the object place recognition test was carried out to examine whether SH mice have a deficit in the hippocampus-dependent memory^34^ (Fig. 1f). Again, SH mice investigated an object in a novel position significantly more than an object in a familiar position, demonstrating that SH mice do not have a hippocampus-dependent memory deficit. GH and SH mice showed comparable body mass changes (Extended Data Fig. 1a). In addition, SH mice showed a comparable basal locomotor activity and anxiety level to those in GH mice (Extended Data Fig. 1b, c).

**Figure 1.**
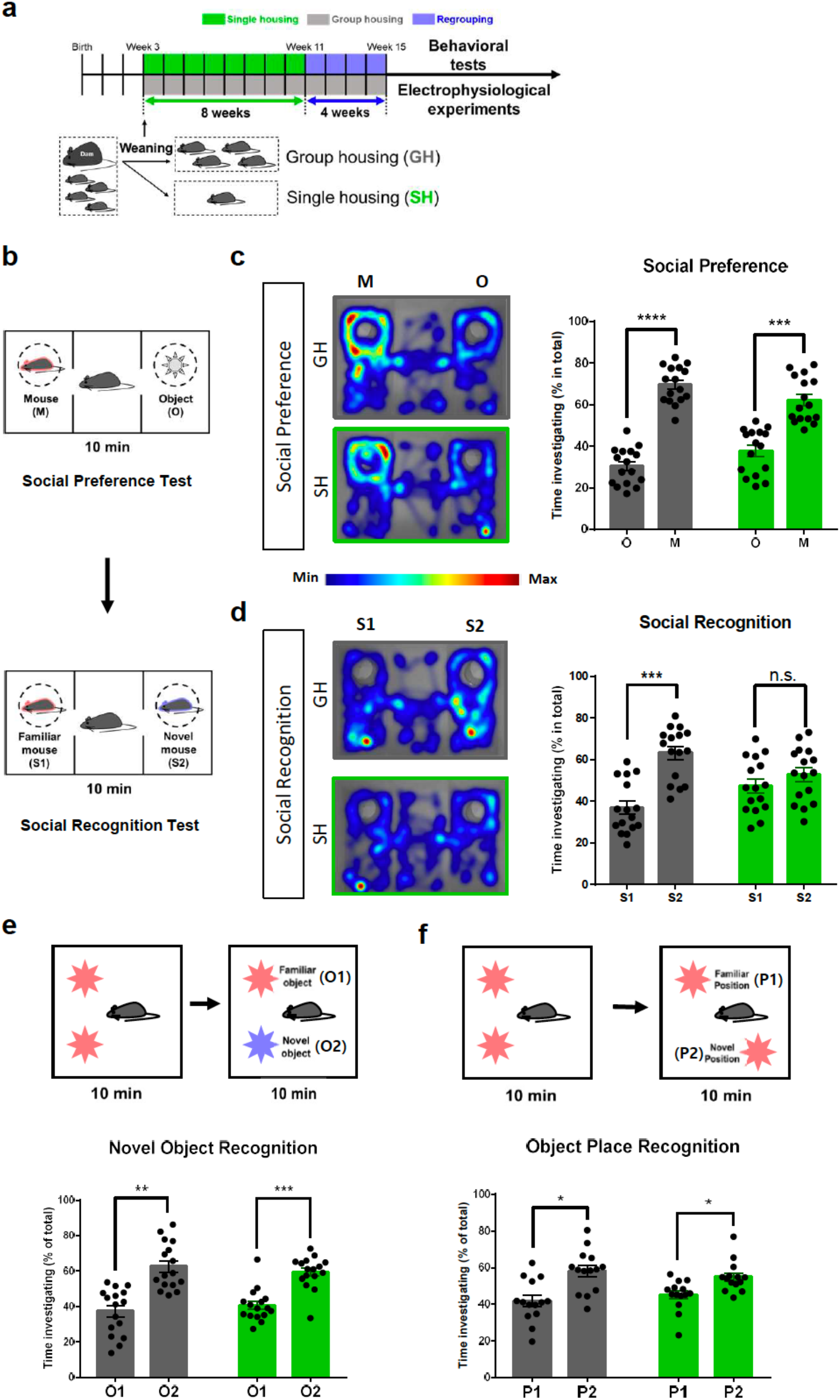
Juvenile social isolation impairs social recognition in adulthood, even after re-socialization. **a,** Timeline of social isolation model used in this study. Single housed (SH) mice are singly housed for 8 weeks and regrouped for at least 4 weeks before tests. Group housed (GH) mice are compared to SH mice as a control group. **b,** Schematic diagram for three-chamber social behavior tests. M, target conspecific; O, inanimate object; S1, familiar conspecific; S2, novel conspecific. **c,** Left: Representative heatmap images of social behavior during the social preference test. Right: In the social preference test, both GH and SH mice with the resocialization period spent significantly more time exploring a target conspecific than an inanimate object. Two-tailed paired t-test: GH, \*\*\*\**P* < 0.001, n = 16 biologically independent mice; SH mice, \*\*\**P* = 0.0004, n = 16 biologically independent mice. **d,** Left: Representative heatmap image of social behavior during the social recognition test. Right: In the social recognition test, SH mice did not show a significant preference towards a novel conspecific versus a familiar conspecific, while GH mice shows a significant preference for a novel conspecific. Two-tailed paired t-test: GH, \*\*\**P* = 0.0008, n = 16 biologically independent mice; SH mice, *P* = 0.4139, n = 16 mice. n.s., not significant. **e,** Top: Schematic diagram for novel object recognition test. O1, familiar object; O2, novel object. Bottom: Both SH mice and GH mice spent significantly more time investigating the novel object than the familiar object Two-tailed paired t-test: GH, \*\**P* = 0.0015, n = 16 biologically independent mice; SH mice, \*\*\**P* = 0.0009, n = 16 mice. **f,** Top: Schematic diagram for object place recognition test. P1, familiar position; P2 novel position. Bottom: Both SH mice and GH mice spent significantly more time investigating an object with a novel position than the other object with the original position. Two-tailed paired t-test: GH, \**P* = 0.0205, n = 14 biologically independent mice; SH mice, \**P* = 0.0464, n = 14 mice. Data are represented as mean ± SEM.

We also tested the effects of different durations of social isolation and resocialization on social behaviors. When mice were single housed for 2 weeks immediately after weaning, the known critical developmental period for the medial prefrontal cortex (mPFC), and regrouped for 4 weeks, SH mice showed a comparable performance to GH mice in both the social preference and social recognition test (Extended Data Fig. 2a, b). We also tested whether regrouping SH mice for longer than 4 weeks may rescue the impaired social recognition ability in SH mice (Extended Data Fig. 2c). Interestingly, SH mice still showed a deficit in social recognition, not in social preference, even after 8 weeks of regrouping (Extended Data Fig. 2d).

### Single housed mice showed reduced NAcSh-projecting IL neuronal excitability

Previous studies showed that the hippocampus plays a critical role in social recognition^35, 36^. In particular, the connection between the ventral hippocampus CA1 to the nucleus accumbens shell (NAcSh) was shown to be important for distinguishing a novel conspecific^36^. As the mPFC is known to regulate social behavior via its connection to subcortical regions, including the NAcSh, which is also implicated in social behaviors^37, 38, 39^, we hypothesized that the neural projection from the mPFC to the NAcSh might be involved in social recognition. Neurons in the ventral mPFC regions, including the deep layer of the infralimbic cortex (IL), were heavily labelled by retrograde green fluorescent protein (GFP) virus injection into the NAcSh (Fig. 2a, Extended Data Fig. 3). Although there was also the neuronal projection from the prelimbic (PL) to the NAcSh, the neuronal density was far less compared to the projection from the IL to the NAcSh, which was consistent with previous reports^40, 41^ (Fig. 2b). In *ex vivo* brain slice wholecell patch clamp recordings, NAcSh-projecting IL neurons in the deep layer exhibited significantly reduced neuronal excitability in SH mice compared to in GH mice (Fig. 2c). Reduced neuronal excitability was not observed by NAcSh-projecting PL neurons (Fig. 2d), showing that the impacts of single housing on neuronal excitability is cell-type specific. The excitatory synaptic drive (excitatory postsynaptic current, EPSC) onto NAcSh-projecting IL neurons of SH mice and other electrophysiological properties were comparable to those of GH mice (Extended Data Fig. 4).

**Figure 2.**
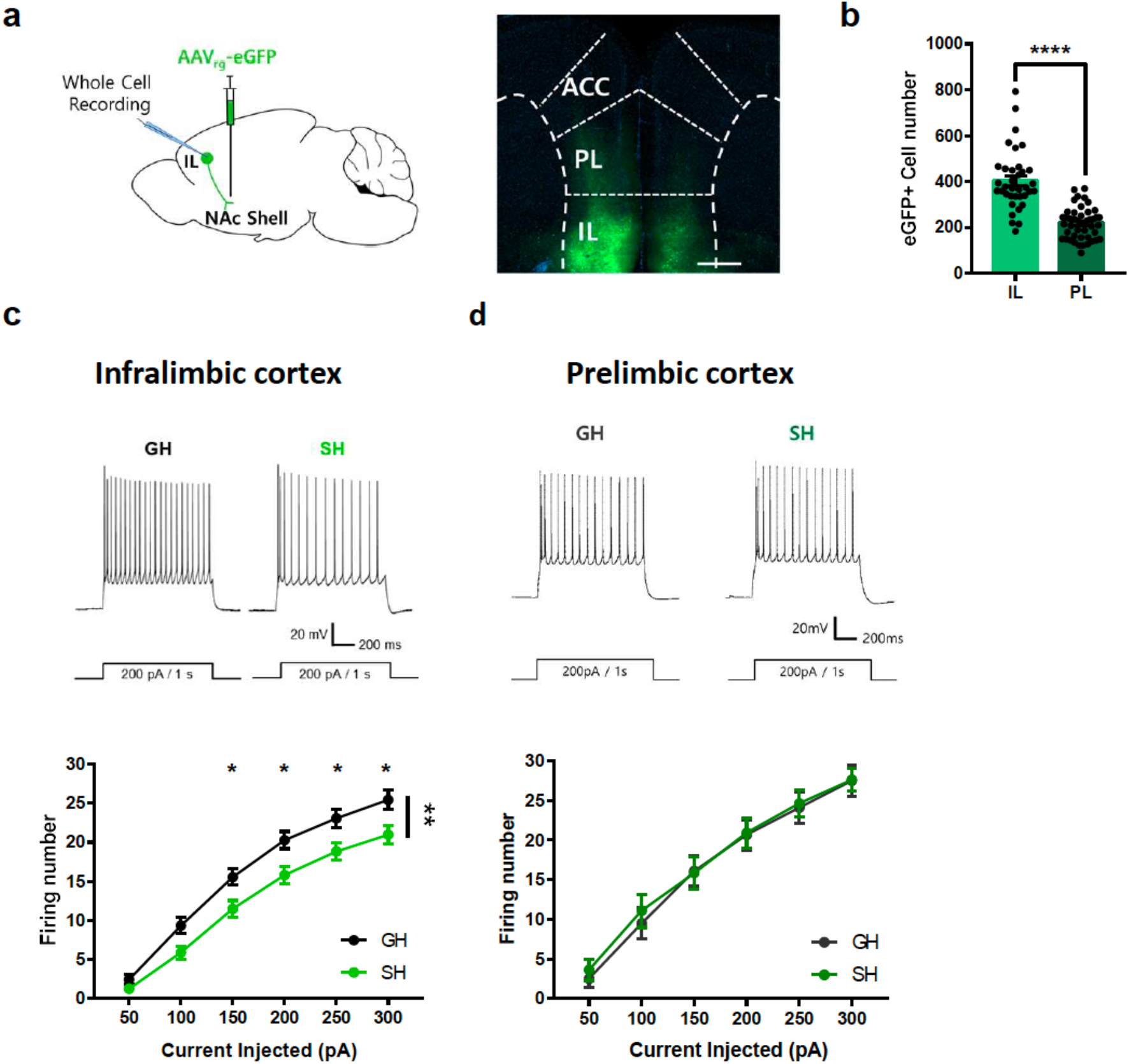
Excitability of NAcSh-projecting IL neuron is decreased in SH mice. **a**, Left: Strategy to label nucleus accumbens shell (NAcSh)-targeting IL neurons by injecting a retrograde adeno-associated virus (AAVrg) expressing eGFP into the NAcSh. Right: Representative image showing eGFP expression in the prefrontal cortex. Scale bar, 500 μm. **b,** Quantification of eGFP-labeled cells in the infralimbic (IL) and prelimbic (PL) cortex. Twotailed unpaired t-test, ****P < 0.0001. IL, 47 slices from 16 mice; PL, 36 slices from 15 mice. **c,** Top: Representative traces of whole cell patch clamp recording in NAcSh-projecting IL neurons. Step currents (200 pA, 1 s) were given. Bottom: Summary data of the number of action potentials evoked in response to currents steps. The number of spikes were significantly decreased in ILà NAcSh neurons in SH mice compared to GH mice. Two-way repeated measure ANOVA, effect of housing, *F*_1, 83_ = 8.806, \*\**P* = 0.0039, n = 37 cells from 14 mice (GH), 49 cells from 13 mice (SH). Sidak’s multiple comparisons test: 150 pA, \**P* = 0.0308; 200 pA, \**P* = 0.0131; 250 pA, \**P* = 0.0220. **d,** Top: Representative traces of whole cell patch clamp recording in NAcSh-projecting PL neurons. Bottom: Summary data of the number of action potentials evoked in response to 300 pA currents steps. NAcSh-projecting PL neurons in SH and GH mice showed comparable excitability. Two-way repeated measure ANOVA, effect of housing, *F*_1, 23_ = 0.05851, *P* = 0.8110, n = 12 cells from 6 mice (GH), 13 cells from 4 mice (SH).

### NAcSh-projecting IL neurons are activated by a familiar conspecific

To confirm the social recognition deficit in SH mice in a different behavioral paradigm, we used social habituation/recognition tasks (Fig. 3a). During the habituation session, mice were consecutively exposed to the same target mouse. Both GH and SH mice showed significantly decreased interacting time with the conspecifics in the habituation session (Fig. 3b). In the social recognition session, which was carried on the next day, mice were allowed to explore either an empty cup, a novel conspecific, or a familiar conspecific. GH mice showed a significant preference toward the novel mouse over either the empty cup or the familiar mouse (Fig. 3c). However, SH mice spent comparable time exploring the novel and familiar conspecific (Fig. 3c), showing that SH mice are impaired in social recognition.

**Figure 3.**
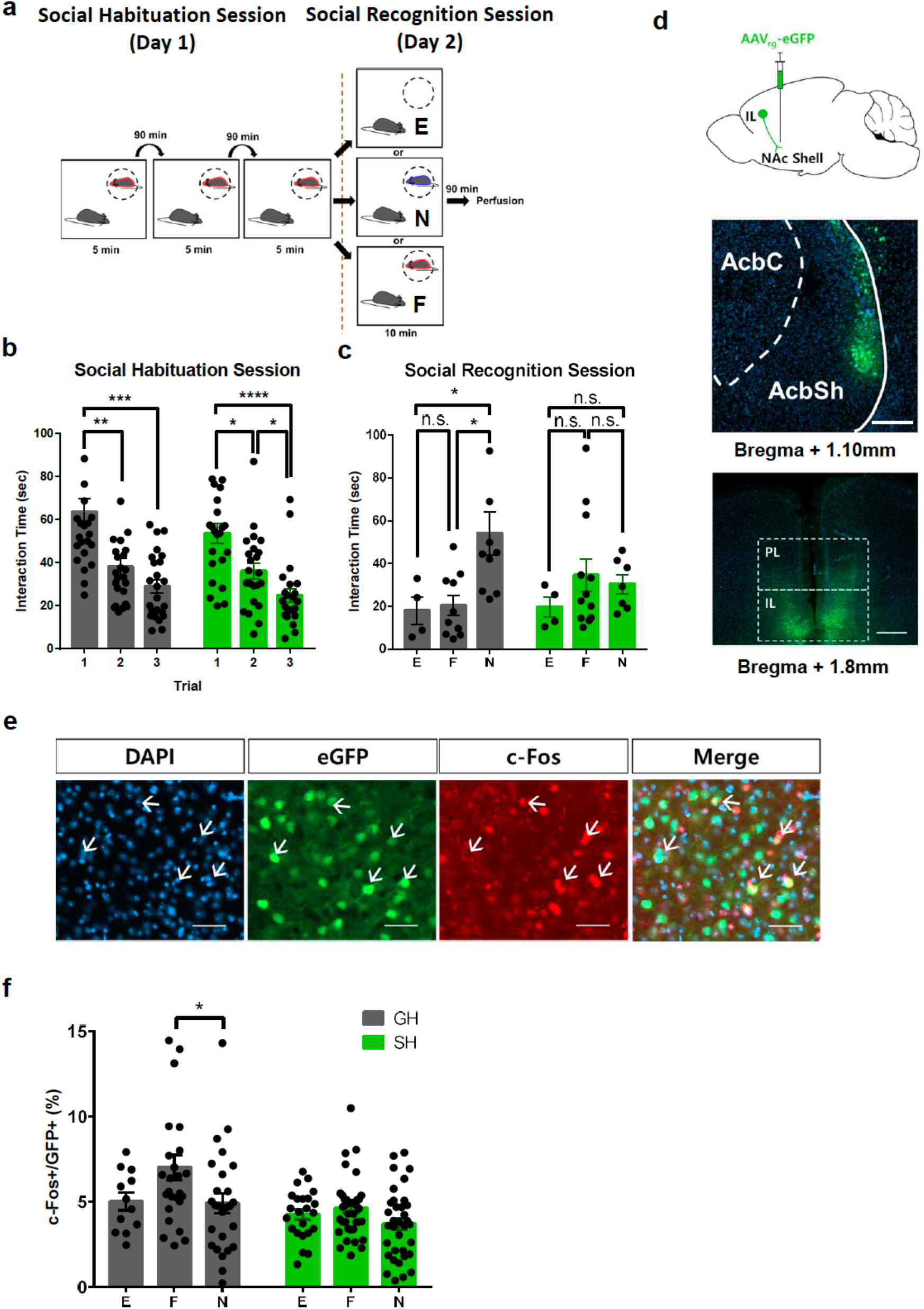
NAcSh-projecting IL neurons are activated by encountering a familiar mouse. **a,** Experimental scheme for the social habituation/social recognition task. Habituation session: GH or SH mice were exposed to the same conspecifics three times for 5 minutes with a 90-minute interval. Social recognition session: subject mice were exposed to familiar conspecifics, novel conspecifics, or an empty cup. E, empty cup; F, familiar target conspecific; N novel target conspecific. **b,** Both GH and SH mice show significantly decreased interaction times during the social habituation session. One-way repeated measure ANOVA followed by Tukey’s multiple comparison tests: GH, 1 versus 2 *t*_23_ = 5.642, \*\**P* = 0.0016; 1 versus 3 *t*_23_ = 6.882, \*\*\**P* = 0.0002; 2 versus 3 *t*_23_ = 2.595; SH, 1 versus 2 *t*_22_ = 4.074, \**P* = 0.0227; 1 versus 3 *t*_22_ = 8.233, \*\*\*\**P* < 0.0001; 2 versus 3 *<u>t</u>*_22_ = 4.031, \**P* = 0.0045. n = 24 mice (GH), 23 mice (SH). **c,** SH mice failed to discriminate familiar conspecifics from novel conspecifics. GH mice showed significantly higher interacting time with novel conspecifics than with familiar conspecifics. One way ANOVA followed by Tukey’s multiple comparison tests: GH, E versus F *q*_21_ = 0.2487; E versus N *q*_21_ = 3.672, \**P* = 0.0427; F versus N *q*_21_ = 4.775, \**P* = 0.0114; SH, E versus F *q*_20_ = 1.722; E versus N *q*_20_ = 1.13, F versus N *q*_20_ = 0.6017. n = 4 mice (GH-E), 10 mice (GH, F), 10 mice (GH, N), 4 mice (SH, E), 12 (SH, F), 7 (SH, N). **d,** Top: Strategy to label NAcSh-projecting IL neurons by injecting the retrograde fluorescence virus (AAVrg-eGFP) in the NAcSh. Middle: Representative image of the brain slices of NAcSh where the virus was injected. Scale bar, 200 μm. Bottom: Representative image of mPFC expressing retrogradely transported eGFP virus. Scale bar, 500 μm. **e,** Representative images showing fluorescence for eGFP-labeled NAcSh-projecting IL neurons (greed), c-Fos (red), DAPI (blue), and merged image. White arrows indicate co-labeled cells for eGFP and c-Fos. Scale bar, 20 μm. **f,** GH mice shows significantly increased number of c-Fos-positive NAcSh-projecting IL neurons when encountered by familiar conspecifics. However, SH mice show no difference in the number of activated neurons depending on the familiarity of the target mice. One-way ANOVA followed by Tukey’s multiple comparisons test: GH, *F*_2,61_ = 3.348, \**P* = 0.0417, Tukey’s multiple comparisons test: E versus F *q*_61_ = 2.578,; E versus N *q*_61_ = 0.13; F versus N *q*_61_ = 3.408, \**P* = 0.0491. n = 12 slices from 4 mice (GH, E), 26 slices from 10 mice (GH, F), 26 slices from 10 mice (GH, N); SH, E versus F *q*_93_ = 1.079; E versus N *q*_93_ = 1.662; F versus N *q*_93_ = 3.121. n = 23 slices from 4 mice (SH, E), 38 slices from 12 mice (SH, F), 35 slices from 7 mice (SH, N).

Since the subject mice were exposed to either a familiar or a novel mouse in this behavioral paradigm, we could examine whether the NAcSh-projecting mPFC IL neurons are differentially activated by distinct social stimuli (familiar vs. novel conspecific) by monitoring the neuronal activity using c-Fos immunohistochemistry after the social recognition session (Fig. 3d). A recent study showed that mPFC IL neurons are activated either by a familiar or by a novel mouse^42^. To monitor the neuronal activity selectively in the NAcSh-projecting IL neurons after social interaction either with a familiar or a novel mouse, we injected a retrograde adeno-associated virus (AAVrg) encoding the enhanced green fluorescent protein (eGFP) into the NAcSh in both SH and GH mice (Fig. 3d). When we analyzed c-Fos and eGFP co-labelled neurons in the IL after the mice were exposed to either a novel-or familiar conspecific, intriguingly, GH mice, who interacted with a familiar conspecific, showed significantly more co-labelled neurons in the IL than GH mice who interacted with a novel mouse (Fig. 3e). Furthermore, the SH mice did not show this familiar conspecific-induced c-Fos activation (Fig. 3e). These results suggest that NAcSh-projecting IL neurons are activated by the consequence of interacting with familiar conspecifics.

### IL-NAcSh circuit is required for social recognition

To examine whether the NAcSh-projecting IL neurons are critically involved in social recognition, we chemogenetically inhibited these neurons in normal GH mice (Fig. 4a). The hM4Di receptor is specifically expressed in the NAcSh-projecting IL neurons (Fig. 4b, Extended Data Fig. 3b). hM4Di or control virus (mCherry)-expressing mice were intraperitoneally (i.p.) injected with clozapine-N-oxide (CNO), which was confirmed to reduce neuronal excitability in hM4Di-expressing neurons (Fig. 4c, d). Behavior tests were carried out 40 minutes after the CNO or vehicle injection. Inhibiting NAcSh-projecting IL neurons did not affect the social preference of the mice (Fig. 4e). Interestingly, when these neurons were inhibited, mice showed a reduced investigation time towards a novel mouse and decreased preference index for social recognition compared to vehicle-injected mice (Fig. 4f). CNO injection did not affect either social preference or recognition in control virus-expressing mice (Fig. 4e, f). These results demonstrate that the NAcSh-projecting IL neuronal activity is required for normal social recognition. Importantly, inhibiting NAcSh-projecting IL neurons did not affect the novel object recognition or object place recognition test (Extended Data Fig. 5a, b). In addition, inhibiting this neuronal activity did not affect the basal locomotor activity or anxiety level (Extended Data Fig. 5c-e). The forced swim test (FST) was also performed to test whether manipulating this neuronal activity may cause any depressive-like behavior^43^. However, inactivating NAcSh-projecting IL neurons did not affect the immobility time in the FST (Extended Data Fig. 5f).

**Figure 4.**
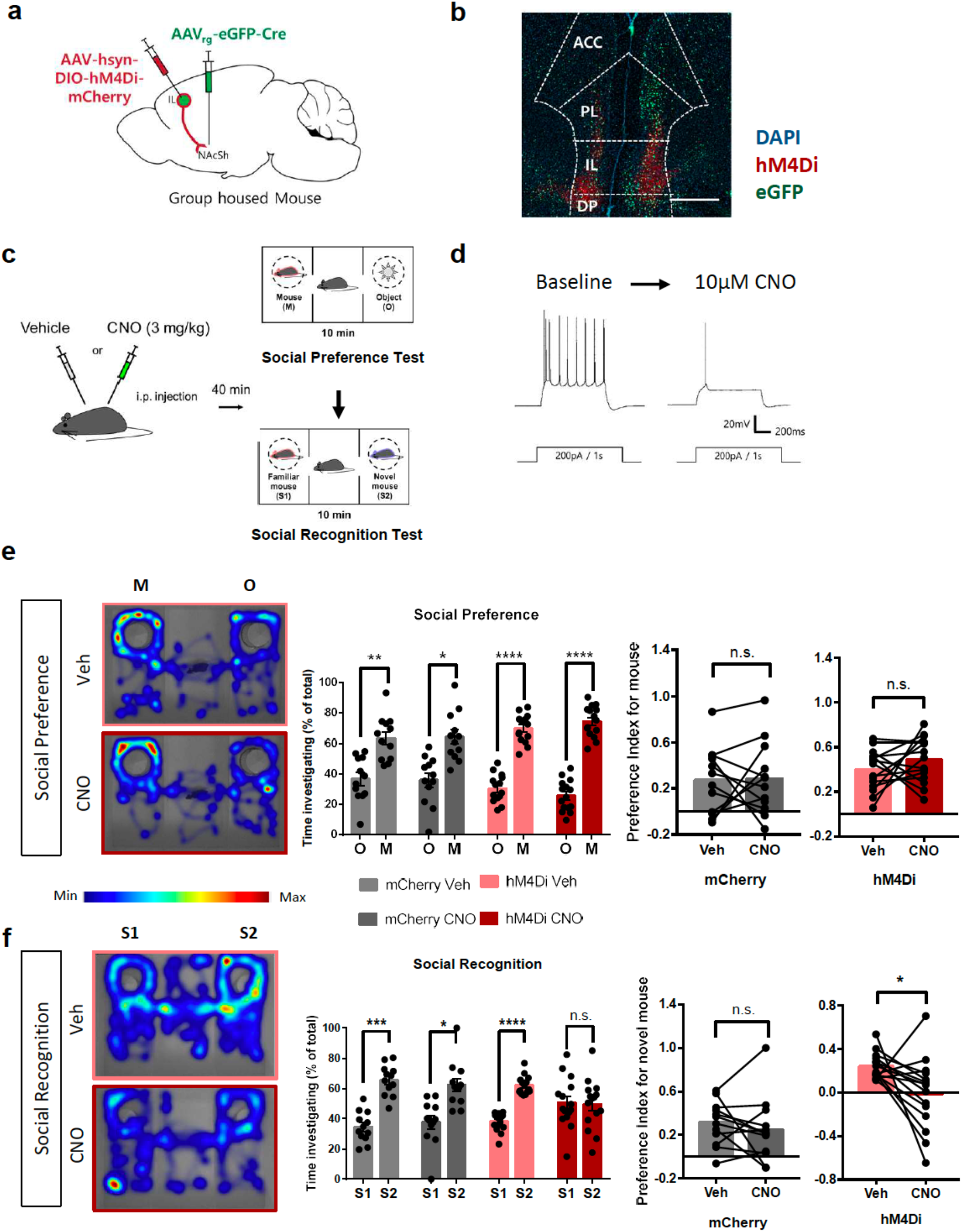
Chemogenetic inactivation of IL-NAcSh neurons selectively impairs social recognition. **a,** Strategy to express Gi-coupled hM4Di receptor or mCherry specifically in NAcSh-projecting IL neurons by injecting a Cre-dependent hM4Di or mCherry virus in the IL and eGFPrg-Cre virus in the NAcSh of group housed mice without social isolation. **b,** Representative image of mPFC expressing hM4Di protein preferentially in IL neurons. Scale bar, 500 μm. **c,** Experimental scheme for inhibiting hM4Di-expressing ILàNAcSh neurons. Mice were injected with either 3 mg/kg clozapine-N-oxide (CNO) or vehicle (saline) and subjected to the 3-chamber tests. **d,** Representative traces of slice whole cell patch clamp recordings confirming the inhibitory effect of CNO in a hM4Di-neuron. **e,** Left: Representative heatmap image of a CNO or vehicle injected hM4Di-expressing mouse during social preference test. Middle: Inactivation of IL-NAcSh neurons did not affect the social preference. Two-tailed paired t-test: mCherry/vehicle, \*\**P*= 0.0081, n =12 mice; mCherry/CNO, \**P* = 0.0104, n = 12 mice; hM4Di/vehicle, \*\*\*\**P* < 0.0001, n = 15 mice; hM4Di/CNO, \*\*\*\**P* < 0.0001, n = 15 mice. Right: Social preference indices before and after CNO treatment were calculated from data shown in the middle panel. Two-tailed paired t-test: mCherry, *P* = 0.8871; hM4Di, *P* = 0.2551. **f,** Left: Representative heatmap image for social recognition test. Middle: Inactivation of IL-NAcSh neurons impairs social recognition. Two-tailed paired t-test: mCherry/vehicle, \*\*\**P* = 0.0002, n = 12 mice; mCherry/CNO, \**P* = 0.0160, n = 12 mice; hM4Di/vehicle, \*\*\*\**P* < 0.0001, n = 15 mice; hM4Di/CNO, *P* = 0.9007, n = 15 mice. Right: Social recognition index was significantly decreased after CNO injection in hM4Di expressing mice. Two-tailed paired t-testL: mCherry, *P* = 0.4478; hM4Di, \**P* = 0.0260.

It is worthy to note that we also found that the normal GH mice did not distinguish a novel mouse from its cage mate when the NAcSh-projecting IL neurons were inhibited (Extended Data Fig. 6a, b). However, this manipulation did not affect the reciprocal social interaction with a novel conspecific, showing that this neuronal activity is less likely to affect sociability itself (Extended Data Fig. 6c). Taken together, our results strongly suggest that the NAcSh-projecting IL neurons may encode social familiarity and contribute to social recognition.

### Increasing NAcSh-projecting IL neuronal activity rescues the social recognition deficit in SH mice

We next examined whether increasing NAcSh-projecting IL neuronal activity could rescue the social recognition deficit in SH mice. To selectively activate these neurons, we expressed the hM3Dq receptor in NAcSh projecting IL neurons and injected CNO (1 mg/kg, i.p.) 40 min before behavioral tests (Fig. 5a-d). The SH mice showed social preference comparable to GH mice regardless of CNO treatment (Fig. 5e). Interestingly, CNO-injected SH mice spent a significantly longer time exploring the novel mouse, demonstrating that the social recognition deficit in SH mice can be rescued by enhancing the activity of NAc-projecting IL neurons (Fig. 5f).

**Figure 5.**
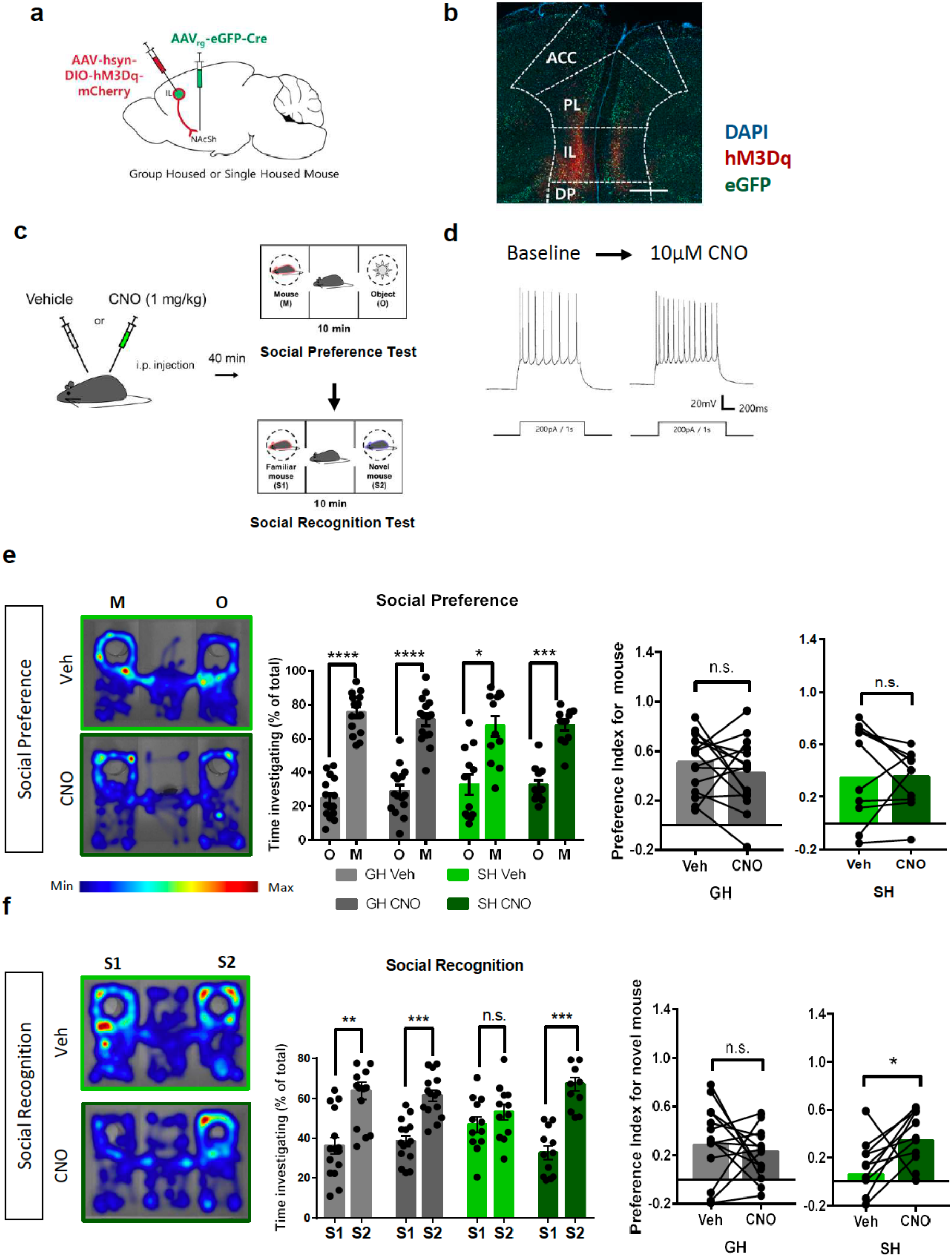
Chemogenetic activation of NAcSh-projecting IL neurons restores the social recognition deficit in SH mice. **a,** Strategy to express Gq-coupled hM3Dq receptor specifically in NAcSh-projecting IL neurons in GH or SH mice. **b,** Representative image of mPFC expressing hM3Dq receptor preferentially in IL neurons. Scale bar, 500 μm. **c,** Strategy to activate hM3Dq expressing ILàNAcSh neurons during 3-chamber social behavior tests by injecting CNO (1mg/kg). **d,** Traces from *ex vivo* whole cell recording validating the activation of hM3Dq receptor by CNO treatment. **e,** Left: Representative heatmap image of a CNO or vehicle-injected hM3Dq-expressing SH mouse during the social preference test. Middle: Activating IL-NAc shell neurons had no significant effect on the social preference in SH and GH mice. Two-tailed paired t-test: GH/vehicle, \*\*\*\**P* < 0.0001, n = 15 mice; GH/CNO, \*\*\*\**P* < 0.0001, n = 15 mice; SH/vehicle, \**P* = 0.0143, n = 12 mice; SH/CNO, \*\*\**P* = 0.0001, n = 12 mice. Right: CNO treatment did not affect social preference index in either group. Two-tailed paired t test: GH, *P* = 0.3171; SH, *P* = 0.9528. **f,** Left: Representative heatmap image for social recognition test. Middle: Inactivation of IL-NAcSh neurons rescues social recognition deficit in SH mice. Two-tailed paired t-test: GH/vehicle, \*\**P* = 0.0064, n = 15 mice; GH/CNO, \*\*\**P* = 0.0009, n = 15 mice; SH/vehicle, *P* = 0.4510, n = 12 mice; SH/CNO, \*\*\**P* = 0.0004, n = 12 mice. Right: CNO treatment significantly increases social recognition index in SH mice. Two-tailed paired t-test: GH, *P* = 0.6956; SH, \**P* = 0.0109.

## Discussion

In this study, we revealed the novel function of the IL-NAcSh circuit in social recognition and the behavioral and electrophysiological impacts of jSI on this circuit. Animals have the tendency to seek novelty both in social and nonsocial stimuli to successfully respond to changing environments^44, 45^. Various animal models of ASD and schizophrenia have shown the defects in social recognition, social novelty preference, or social memory^7, 46, 47, 48^, which may reflect the problems in human patients with ASD or schizophrenia in expanding their social relationships.

Social recognition/memory is known to be governed by several brain regions including the mPFC and the hippocampal subregions such as the dorsal CA2 and ventral CA1 (vCA1)^35, 36, 49^. The vCA1-mPFC projection is hyperactive in the *Mecp2* knockout mice, which show a social memory deficit, suggesting that this projection is involved in social memory^50^. A recent study showed that the vCA1 neurons also projecting to the NAcSh are critically involved in social memory, particularly when responding to a familiar mouse^36^. In this study, we found that mPFC IL neurons projecting to the NAcSh are also activated by a familiar mouse in normal mice, suggesting that the IL-NAcSh projection also encodes social familiarity. NAcSh-projecting IL neurons in SH mice were not activated in response to the familiar mouse, which may underlie a social recognition deficit in SH mice. Interestingly, SH mice showed decreasing social interaction time in the habituation session (Fig. 3), suggesting that single housing may impair either the consolidation or expression, not the acquisition, of social memory. Further studies with longitudinal *in vivo* imaging will provide more information on the role of the IL-NAcSh circuit in social memory.

It has been shown that dopaminergic projections from the ventral tegmental area (VTA) are specifically involved in preference for social novelty, not for object novelty^51, 52^. The importance of the IL-NAcSh circuit was initially announced regarding drug seeking behavior and addiction. As the nucleus accumbens is the key node for the mesolimbic dopaminergic pathway, neuronal projection towards the NAcSh was studied in the perspective of the reward and motivation system^53, 54, 55, 56^. Social isolation is also known as a factor contributing to impairment of motivation and drug seeking behavior. For example, socially isolated mice are significantly more susceptible to alcohol consumption^57^. SH mice in our study exhibited the defect in social recognition without the impairment in novel object recognition, implying the likelihood of an alteration in the mesolimbic dopaminergic pathway encompassing the IL-NAcSh circuit by jSI. It has been shown that chronic social isolation in adult rats decreased CREB activity as well as the excitability of the NAcSh neurons, which may underly depression-like behaviors in socially isolated rats^58^. We found that the excitability of NAcSh-projecting IL neurons is significantly decreased in SH mice, showing that the excitability of both mPFC and NAcSh neurons are affected by social isolation. Consistently, it has been shown that jSI reduced the excitability of a subtype of mPFC pyramidal neurons in layer 5^59^. In addition, a recent study showed that 2 weeks of social isolation followed by 4 weeks re-grouping impairs social interaction and reduces the excitability of mPFC neurons projecting to pPVT^19^. Together, these results suggest that neuronal subpopulations in the mPFC serve unique social functions (e.g. social interaction and recognition) through projecting to distinct subcortical areas. The molecular mechanism underlying reduced excitability in NAcSh-projecting IL neurons remains to be investigated. It would be interesting to examine the role of CREB in our jSI model.

Previous studies using jSI model reported that first 2 weeks after weaning (P21-35) is the critical period for normal social preference^18, 20, 59, 60^. However, we found that this period was not long enough to induce the long-lasting deficit in social recognition (Extended Data Fig. 2a, b). Piskorowski et al. showed that schizophrenia model mice (Df16(A)^+/-^) exhibited defects in social recognition and its related alterations of neuronal properties occurred only in their adult phase, after postnatal 5 weeks (P35), implying that the postnatal development for normal social recognition could be completed later than that for normal social preference^49^. Our data indicates that the critical period for social recognition is within postnatal 11 weeks. However, it is worthy to note that the impacts of social isolation and re-socialization on social behaviors can be sensitive to the experimental conditions such as the duration of isolation/re-grouping and the mode of re-socialization^18, 30, 60^.

Although the deficits in social preference or sociability has been extensively investigated in animal models of ASD, a lack of differentiation between individuals has also often been reported both in ASD patients and animal models^7, 61, 62, 63^. Our study will contribute to understand not only the biological mechanism for social recognition, but also the pathophysiology underlying social recognition impairments in ASD and related disorders.

## Methods (Online)

### Animals

Male C57BL/6NHsd mice were purchased from Koatech (South Korea). Purchased mice were introduced to the animal facility at Seoul National University College of Medicine on postnatal day 14 (P14) with their dam. Four mice were allocated into a standard mouse cage with their dam. At P21, mice were weaned from their dam. From P70 to P90, the mice may experience the surgical procedure depending on the experimental needs. Social conspecifics C57BL/6NCrljOri were purchased by another breeder in Korea (Orient Bio) at P35 and used for the social behavior test after an adjustment period of 1 week. All mice were given by a fixed 12 hours light-dark cycle (light on: 8:00-20:00; light off: 20:00 – Next day 8:00). Food and water were provided *ad libitum*. Experiments were conducted in accordance with the Seoul National University, College of Medicine, Department of Physiology and approved by the Institutional Animal Care and Use Committee of Seoul National University (IACUC #: SNU 190501-5-2).

### Social isolation

At P21, group-housed (GH) mice were removed from their dam, but still housed with their littermates. Single-housed (SH) mice were individually housed in the standard mouse cage from P21 to P77 (isolated for 8 weeks) or from P21 to P35 (for 2 weeks). After 8 weeks of isolation, single isolated mice were re-grouped with their littermates.

### Surgical procedures

Mice were anesthetized with a Zoletil (30 mg/kg) and Rompun (10 mg/kg) mixture prior to the surgical procedure by i.p. injection with the injection volume varying depend on body weight. The anesthetized mouse was set on the rodent stereotaxic apparatus. A heating pad was placed below the mouse to prevent rodent hypothermia during the stereotaxic surgery. An appropriate region of the skull was drilled with a stereotaxically fastened hand drill assembled with a 0.4 mm drill bit. Virus injection was performed bilaterally. The infralimbic cortex (AP +2.4 from bregma, ML ±1.0 from midline, DV −2.3 from skull; 10° angle given to avoid virus expression in the prelimbic cortex) and nucleus accumbens shell (AP +1.94 from bregma, ML ± 0.55 from midline, DV −4.15 from skull) were targeted for virus injection. A glass capillary (#504949, WPI, USA) is pulled out sharply using the Dual-Stage Glass Micropipette Puller (NARISHIGE, USA). Virus information is summarized below. The virus was diluted with filtered ACSF to the desired titer. Nanoliter 2010 (WPI) or Spritzer Pressure System IIe (Toohey Company, USA) was used for the virus injection. The virus was given towards the target of interest with a speed of 20 uL/min, giving the desired volume of virus depending on the experiment and brain regions. The capillary was set still within the brain at least 5 min before a slow withdrawal. The glass capillary was pulled off slowly and the incised skin was sutured with sterilized suture thread. Animals were rested at least 3 weeks before further experiments.

### Virus information

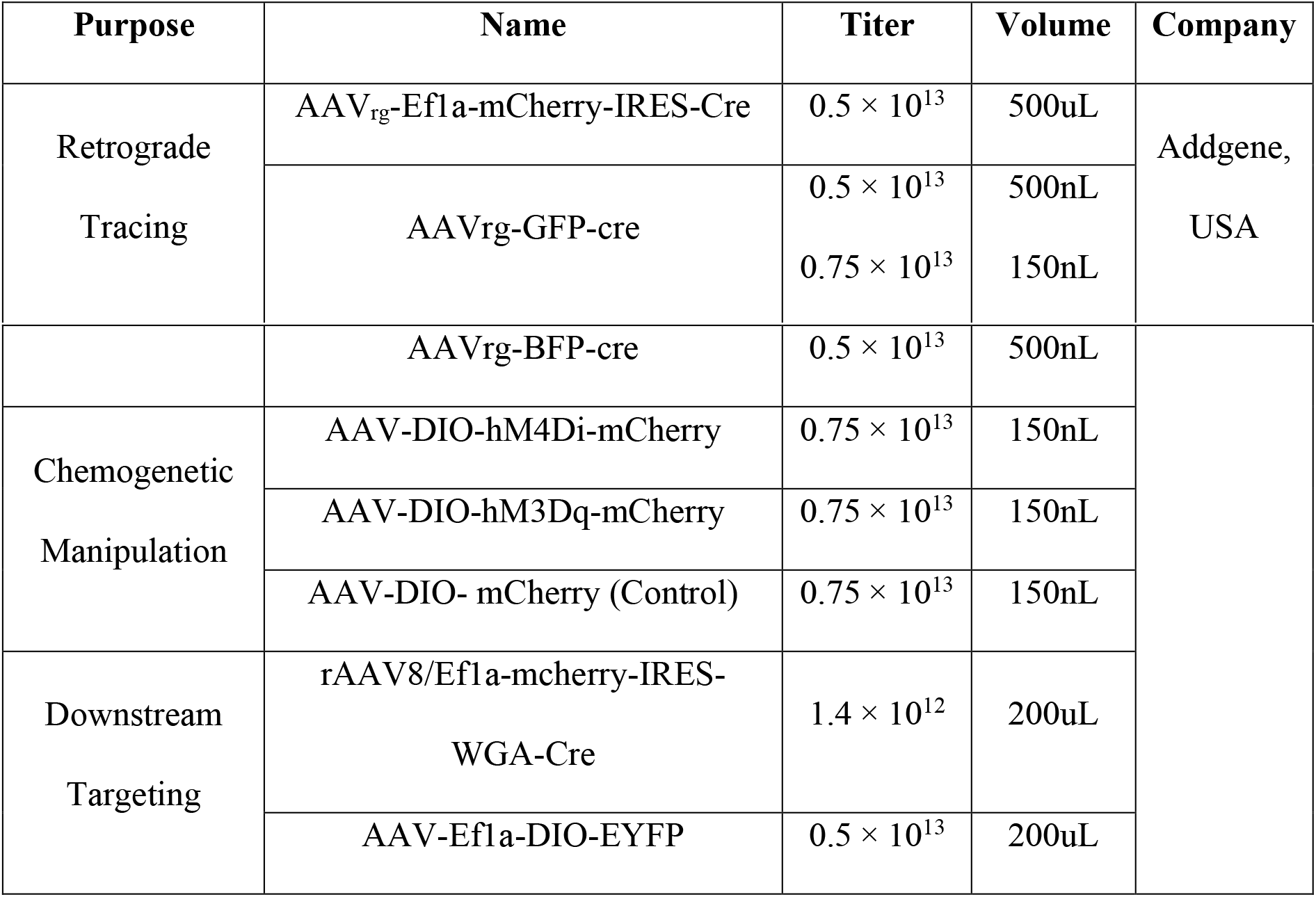

### CNO administration

To activate designer receptors exclusively activated by designer drugs (DREADD), we used clozapine N-oxide (CNO) dihydrochloride (water soluble form, Hello Bio) dissolved in 0.9% normal saline to yield a final concentration of 1 mg/mL. CNO solution (final concentration in the body: 3 mg/kg for hM4Di neuronal inhibition; 1 mg/kg for hM3Dq neuronal activation) or the same volume of saline vehicle was i.p. injected into subject mice 40 minutes before behavioral assessments. CNO or vehicle injection into each mouse was separated by 2 days and counterbalanced for order.

### Behavior Tests

All behavior tests described below were performed in a soundproof chamber with a dim light illumination during the light-on cycle. All behavior tests were recorded by the camcorder and the experimenters were blinded to the experimental conditions. Both the subject and target mice were habituated for hand approach made by a human prior to any behavior test.

### Three-chamber social behavior test

The three-chamber social behavior test was performed as previously described^4^. We used a white opaque acrylic three-chambered apparatuses with two doors between the chambers (40 x 60 cm size, each chamber 20-cm wide). Before the test, target mice were habituated in wire containers placed in side-chambers in the apparatuses for 20 minutes on 2 consecutive days. Each subject mouse was placed in a test consisting of three sessions: (1) habituation, (2) social preference test, (3) social recognition test. For the habituation, the subject mouse was put in the apparatus for 10 minutes with the doors open. After the habituation, the social preference test was performed. The first target mouse was put in a wire container in either the left or right chamber. An inanimate object (mouse-shaped plastic toy) was put in the wire cage in the other side chamber. For the social recognition test, an inanimate object was replaced by another target mouse. During the intervals between sessions, the subject mouse was gently guided to the center room of the apparatus and the doors of the apparatus were closed. After the placement of target mice, the doors were re-opened, and the subject mouse was re-allowed to move freely for 10 minutes. The movements of subject mice during the test were tracked by a mouse tracking software (EthoVision XT 11.5, Noldus). To normalize the variation in the exploration time of each subject, all data were represented by the percentage of time spent investigating one side out of total investigation time for both sides. Preference index (PI) was calculated by following equations (ETM: exploration time for a mouse; ETO: exploration time for an object; ETN: exploration time for a novel mouse (the second target mouse); ETF: exploration time for a familiar mouse (the first target mouse)).

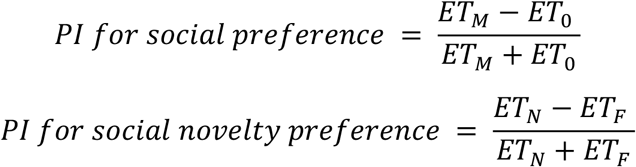

### Reciprocal social interaction test

A clean mouse homecage with a thin layer of bedding was used for the reciprocal social interaction test. Two mice (either two subject mice or one subject mouse and target mouse) were placed in the homecage simultaneously. Mouse interaction is recorded and manually analyzed manually. For the interaction, nose to nose sniffing time and nose to anogenital sniffing time were counted.

### Target familiarization test

Subject mice were allocated to the homecage alone one hour before the test. Subject mice were placed in the white opaque acrylic box (40 cm cubic sized box) with the wired container. The target familiarization test was composed of three session, (1) habituation, (2) familiarization, and (3) retrieval. The target familiarization test took 2 days; habituation and familiarization were done on day 1 while interaction was done on day 2. The habituation session allowed mice to explore the while opaque acrylic box with a wired container for 10 minutes. The target conspecific was introduced into the wired container for 5 minutes. This familiarization was repeated twice more with a 90-minute interval. Subject mice were returned to their homecage after the familiarization. On day 2, the subject mice were exposed to the same environment to the familiarization. However, the subject mice randomly explored the wired container filled with novel conspecific, a familiar conspecific, or the container that was empty.

### Open field test

Subject mice were placed in the center of a white opaque acrylic box (33 x 33 cm size) to move freely for 10 minutes. The center zone was set in 20 x 20 cm size. The movements of subject mice were tracked by EthoVision XT 11.5 (Noldus).

### Novel object recognition test

Prior to the test, subject mice were habituated to a test arena (33 x 33 cm size, white opaque acrylic box) for 20 minutes on 2 consecutive days. On the day of the test, training and test sessions were conducted. For the training session, the subject mouse was put in the center of the test arena containing two identical objects. The mouse was allowed to move freely for 10 minutes. After the training session, the mouse was taken out from the test arena, and one of the objects was replaced with a novel object. Then, the mouse was reintroduced into the test arena to move freely for 10 minutes. The exploration time to each object was scored manually by two independent experimenters. To normalize the variation in the exploration time of each subject, all data were represented by the percentage of time spent investigating one side out of the total investigation time for both sides.

### Object place recognition test

Prior to the test, subject mice were habituated to a test arena (33 x 33 cm size, white opaque acrylic box) for 10 minutes twice on the same day. On the day of the test, training and test sessions were conducted. For the training session, the subject mouse was put in the center of the test arena containing two identical objects. To give a spatial cue to the subject mice, a triangle was put on one side of the box. The mouse was allowed to move freely for 10 minutes. After the training session, the mouse was taken out from the test arena, and one of the objects was moved vertically to a novel location. Then, the mouse was reintroduced into the test arena to move freely for 10 minutes. The exploration time to each object was scored manually by two independent experimenters. To normalize the variation in the exploration time of each subject, all data were represented by the percentage of time spent investigating one side out of the total investigation time for both sides.

### Elevated plus maze test

The apparatus consisted of two open arms (29.5 x 5 x 0.5 cm), two perpendicular closed arms (29.5 x 5 x 16 cm), and a center platform (5 x 5 x 0.5 cm). The apparatus was raised 50 cm above the floor. The open arms had a short wall (0.5 cm) to keep animals from falling or escaping, whereas the closed arms had a tall wall (16 cm). The subject mouse was placed in the center of the apparatus (with its head heading toward open arms) and was allowed to move freely for 10 minutes. The exploration time to each arm was scored using EthoVision XT 11.5 (Noldus).

### Immunohistochemistry

Brains of the subject mice were extracted after carrying out transcardial perfusion by PBS solution followed by 4% PFA solution (T&I, Korea). Perfused brains were emerged into 4% PFA for 24 hours and emerged into 30% (w/v) sucrose solution for 48 hours. Processed brains were frozen at −20°C with clear frozen section compound (FSC22; Leica, Germany) and sliced at a 30 μm thickness. Brain slices were stored in CPS solution for the next usage. Brain slices were incubated with 4% (v/v) goat serum in 0.1% (v/v) triton-X100 in PBS for 40 min. Brain slices were then incubated with primary antibodies above a locker at 4°C for 48 hours.

Information on the primary antibodies are listed below. After 48 hrs, brain slices were incubated with an appropriate secondary antibody (Alexa Fluor Secondary Antibodies; ThermoFisher Scientific, USA) on room temperature for 4 hours. Before mounting the slices, DAPI (Cat #62248; Thermo Scientific, USA; 1:10,000) was added for counterstaining. Images were acquired using a confocal microscope FV3000 (Olympus) or Axio Scan Z1 (Zeiss).

### Electrophysiology

Electrophysiological recordings were performed as described previously^64^. Coronal slices of the mPFC (300 μm thick) were obtained by a vibratome (VT1200s, Leica) after isoflurane anesthesia and decapitation. Slices were cut in ice-cold cutting solution containing (in mM): 93 N-Methyl-D-glucamine (NMDG), 2.5 KCl, 10 MgSO_4_, 0.5 CaCl_2_, 1.25 NaH_2_PO_4_, 30 NaHCO_3_, 25 glucose, 10 HEPES, 5 Na ascorbate, 2 thiourea, 3 Na pyruvate, 12 L-acetyl-cysteine, perfused with 95% O_2_ and 5% CO_2_. The slices immediately transferred to the same cutting solution at 32 °C for 10 minutes, and then transferred to artificial cerebrospinal fluid (ACSF) at room temperature containing (in mM): 125 NaCl, 2.5 KCl, 1 MgCl_2_, 2 CaCl_2_, 1.25 NaH_2_PO_4_, 26 NaHCO_3_, 10 glucose, perfused with 95% O_2_ and 5% CO_2_. The slices were recovered in ACSF for 1 hour before the experiment. All recordings were done within 8 hours from recovery. Brain slices were placed in a submerged chamber and perfused with ACSF for at least 10 minutes before recording. All recordings were made at 32 °C. We used recording pipettes (4-5MΩ) filled with (in mM): 135 K-gluconate, 5 KCl, 2 NaCl, 10 HEPES, 0.1 EGTA, 5 Mg ATP, 0.4 Na_3_ GTP and 10 Tris(di) phosphocreatine (pH 7.20 adjusted by KOH). Data were acquired using an EPC-9 patch-clamp amplifier (HEKA Elektronik) and PatchMaster software (HEKA Elektronik) with a 20 kHz sampling rate, and the signals were filtered at 2 kHz. Synaptic current data were analyzed using Mini Analysis (Synaptosoft). Excitability data were analyzed by customized LabView (National Instruments) analysis programs.

### Statistics

A two-way repeated measure analysis of variance (ANOVA) was performed was followed by an appropriate multiple comparisons post-test to analyze membrane excitability data. The social preference, social recognition, object recognition, and object place recognition test were analyzed by a paired two-tailed t-test. Preference indices and other two-group data were compared using an unpaired two-tailed t-test. All data are represented as the mean ± SEM. Graphpad prism 7.0 (Graphpad software) was used for all the statistical analyses and visualization.

## Acknowledgements

This work was supported by grants to Y.-S.L. (NRF-2017M3C7A1026959, NRF-2019R1A4A2001609), S.J.K. (NRF-2018R1A5A2025964, NRF-2017M3C7A1029611), and G.P. (NRF-2018H1A2A1061381) from the National Research Foundation of Korea. Authors thank Myeong Jong Yoo for helping with managing the single housing mouse colony.

## Author contributions

G.P., C.R., S.J.K., and Y.-S.L. conceptualized the research and design the experiments. Y.-S.L. and S.J.K. supervised the research. G.P., C.R., and S.K. performed behavioral analyses. G.P. and C.R. performed whole cell patch clamp recordings and stereotaxic surgeries. G.P., C.R., and Y.-S.L. wrote the manuscript.

