## Supplementary figures for "Social isolation impairs the prefrontal-nucleus accumbens circuit subserving social recognition in mice"

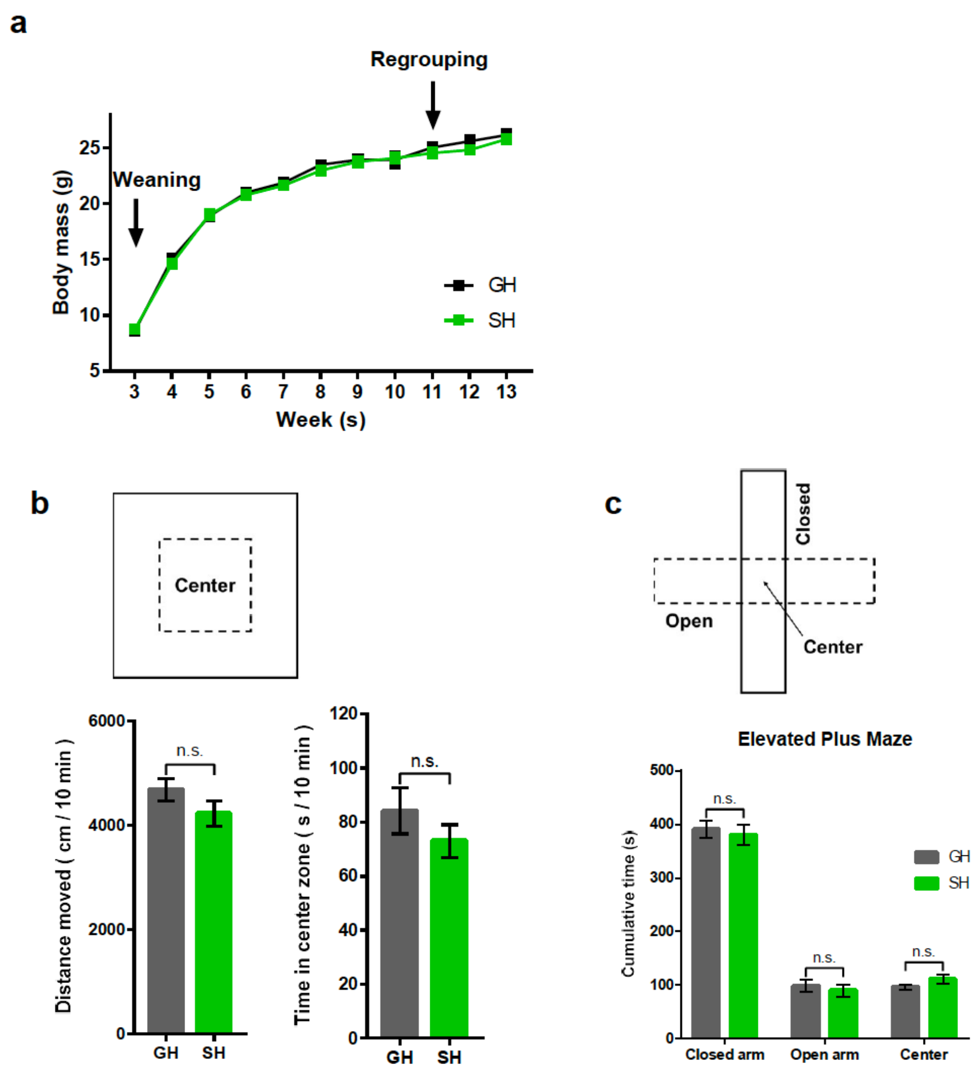


**Extend Data figure 1. Social isolation model with re-socialization selectively affect social recognition in adulthood.**

**a,** Body mass of group housed (GH) and single housed (SH) mice did not show a significant difference before and after the re-socialization period. n = 16 mice (GH), 16 mice (SH).

**b,** Top: Schematic diagram of open field test. Bottom, Left: GH and SH mice showed comparable level of locomotion (n = 8 mice/group, Unpaired two-tailed t-test with Welch's correction, $t$ = 1.425, $P$ = 0.1766). Bottom, Right: time spent in the center zone in the open field test (n = 8 mice/group, Unpaired two-tailed t-test with Welch's correction, $t$ = 1.072, $P$ = 0.3037).

**c,** Top: Schematic diagram of elevated plus maze test. Bottom: SH mice and GH mice showed similar anxiety level (ordinary two-way ANOVA, $F_{2,42}$ = 0.5841, ***$P$ = 0.5621, n = 12 GH mice, n = 12 SH mice [Sidak's multiple comparisons test: Closed arm GH – SH $t_{42}$ = 0.5672; Open arm GH – SH $t_{42}$ = 0.4702; Center GH – SH $t_{42}$ = 0.8024]).

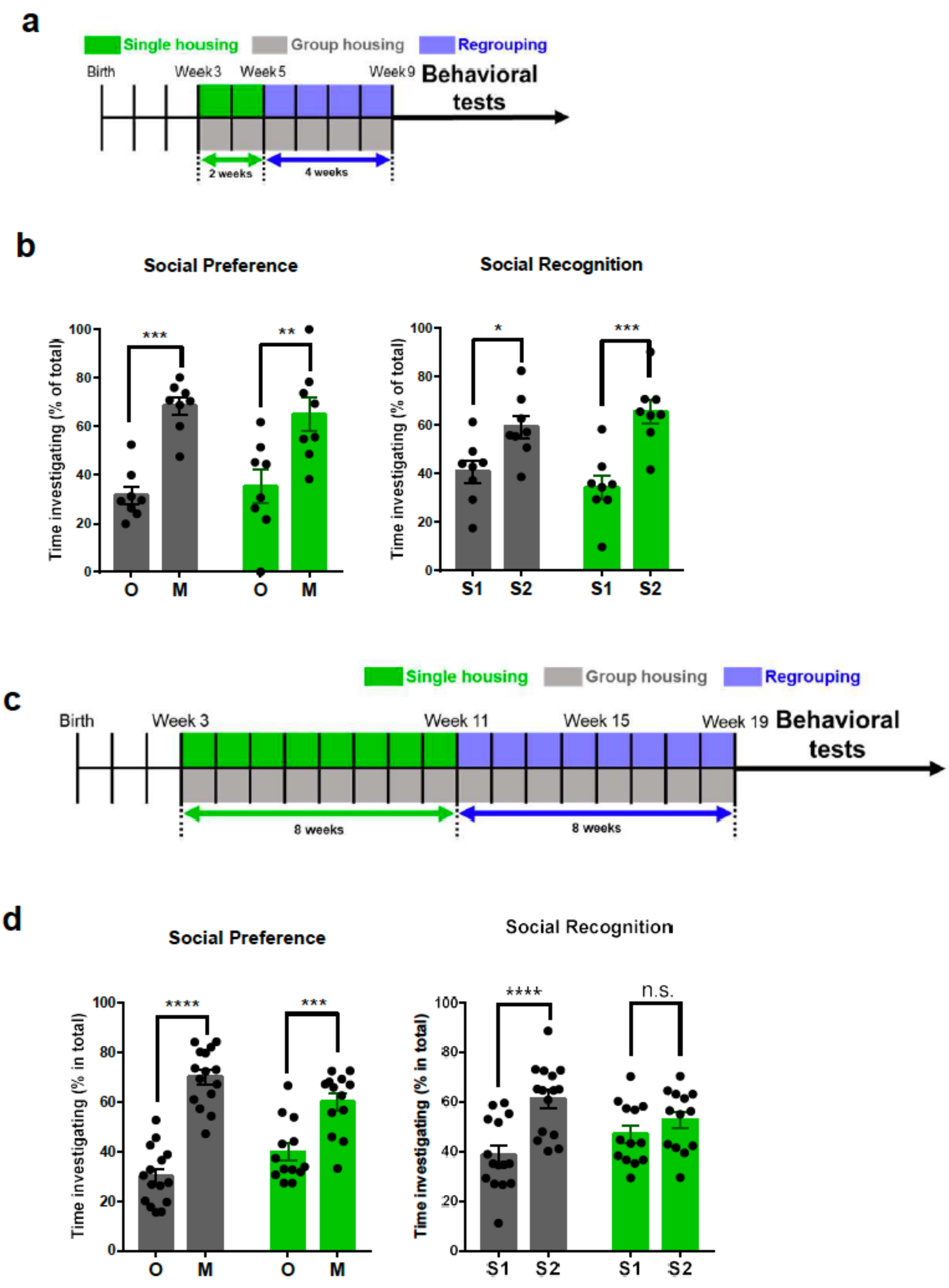


**Extend Data figure 2. Assessments of social behaviors of social isolation models with different experimental timelines.**

**a,** SH mice were isolated for 2 weeks and re-grouped for 4 weeks before behavioral tests. Dark gray color and green color indicate GH and SH mice, respectively.

**b,** Mice singly housed for 2 weeks showed a significant preference towards a conspecifics (n= 8 GH, n= 8 SH, two-way ANOVA, $F_{1, 28} = 0.5225$, $P = 0.5225$ [between-group comparison by Sidak's multiple comparisons test: GH, ***$P = 0.0001$, O - M $t_{28} = 4.72$; SH, **$P = 0.0014$, O – M $t_{28} = 3.804$]). SH mice showed a normal social novelty preference which is similar to the control, GH mice (n= 8 GH, n= 8 SH, two-way ANOVA, $F_{1, 28} = 1.784$, $P = 0.1924$ [between-group comparison by Sidak's multiple comparisons test: GH, *$P = 0.0201$, S1 – S2 $t_{28} = 2.759$; SH, ***$P = 0.0001$, S1 – S2 $t_{28} = 4.647$]).

**c,** SH mice were isolated for 8 weeks and re-grouped for 8 weeks before behavioral tests. Dark gray color and green color indicate GH and SH mice, respectively.

**d,** Mice singly housed for 8 weeks , regrouped for 8 weeks showed a significant preference towards a conspecifics (n= 15 GH, n= 12 SH, ordinary two-way ANOVA, $F_{1, 52} = 9.698$, **$P = 0.0030$ [between-group comparison by Sidak's multiple comparisons test: GH, ****$P < 0.0001$, O - M $t_{52} = 9.252$; SH, ***$P = 0.0001$, O – M $t_{52} = 4.359$]). These single housed mice still showed an impaired social novelty preference compared to the control, GH mice (n= 15 GH, n= 12 SH, ordinary two-way ANOVA, $F_{1, 52} = 5.578$, *$P = 0.0220$ [between-group comparison by Sidak's multiple comparisons test: GH, ****$P < 0.0001$, S1 – S2 $t_{52} = 4.661$; SH, $P = 0.4689$, S1 – S2 $t_{52} = 1.112$]).

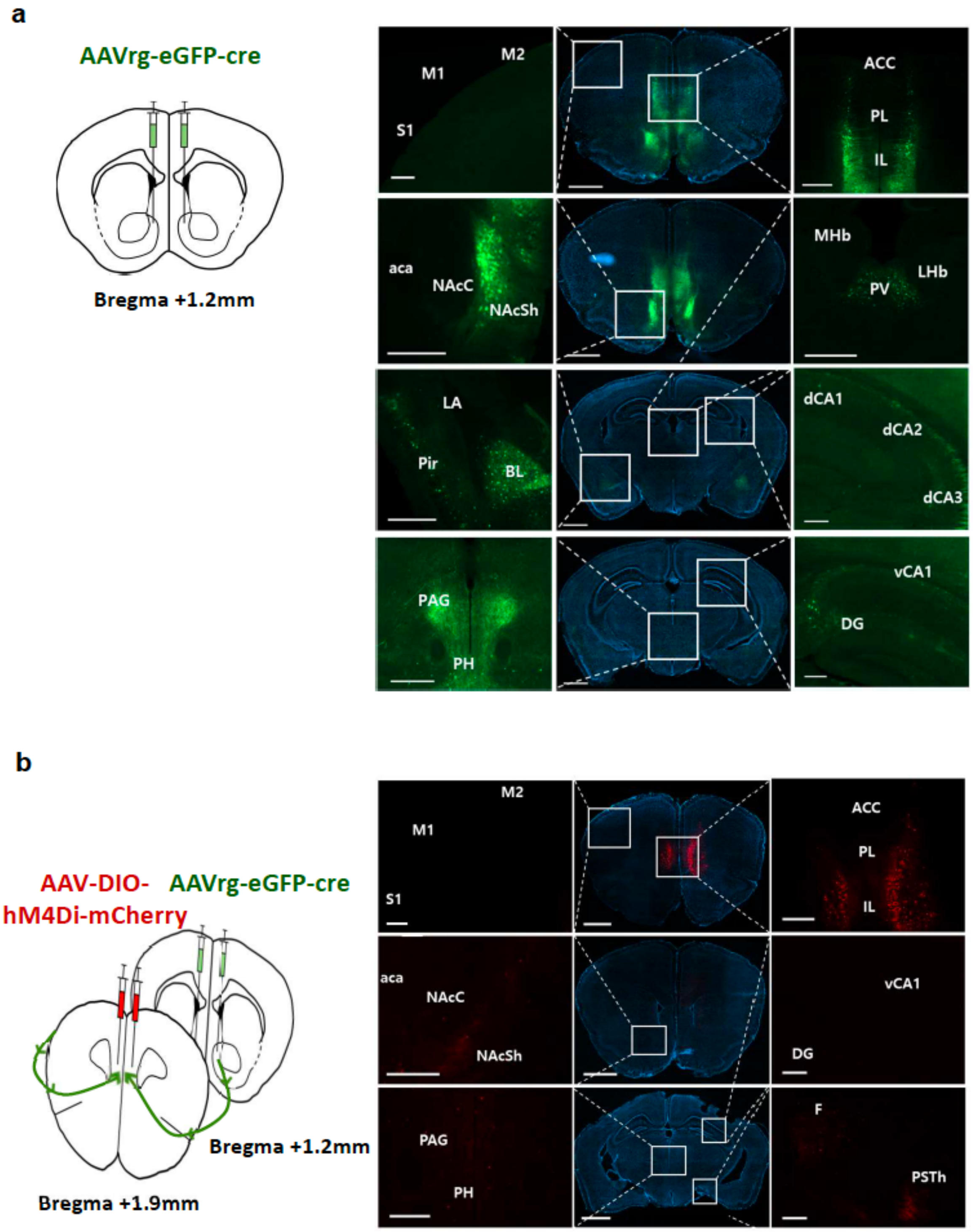


**Extend Data figure 3. NAcSh-projecting IL neurons were specifically labeled.**

**a,** Nucleus accumbens (bregma +1.2mm) shell was targeted to inject a retrograde virus expressing eGFP-Cre. Fluorescence is shown in the mPFC and NAcSh with the highest intensity and in other subcortical regions such as paraventricular nucleus (PV), amygdala (BLA, LA) with observable intensity. Within PFC, infralimbic cortex(IL) projects heavily to NAcSh, compared to prelimbic cortex(PL).

**b,** Infralimbic (bregma +1.9mm) was targeted to inject a cre-dependent hM4Di-mCherry expressing virus. Fluorescence is shown in the PFC, especially in the IL region with its highest intensity, showing its axon terminals in different subcortical regions.

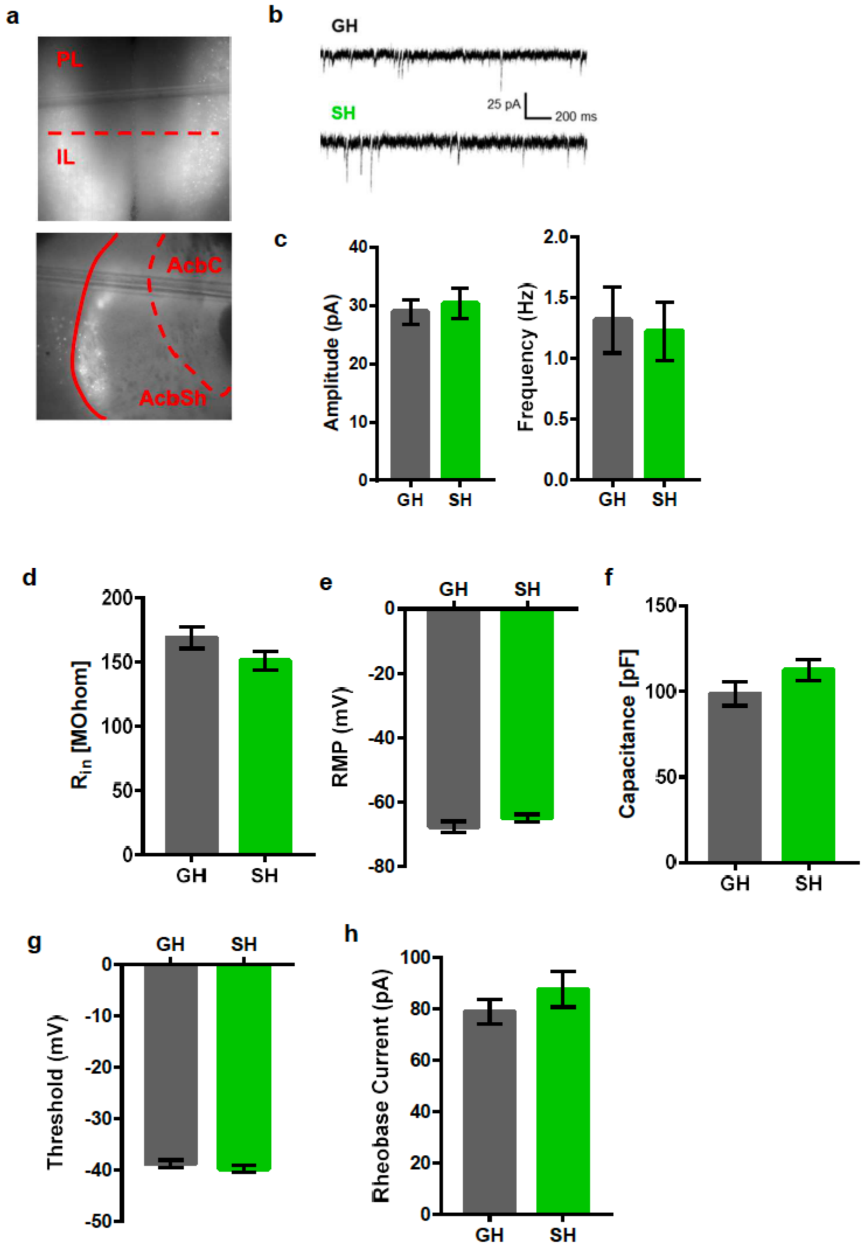


**Extend Data figure 4. SH did not alter excitatory synaptic transmission or basal membrane properties.**

**a,** Representative images of IL and NAcSh showing fluorescence signals from the retrograde GFP virus injected into NAcSh.

**b,** Spontaneous excitatory postsynaptic current (sEPSC) traces of IL→ NAcSh neurons in GH and SH mice.

**c,** Left: sEPSC amplitude of NAcSh-projecting IL neurons in SH mice is comparable to that of GH mice

(Two-tailed unpaired t test with Welch's correction, $t_{29.87}$ = 0.4251, $P$ = 0.6738, n = 14 cells from 2 GH mice, n = 18 cells from 4 SH mice). Right: sEPSC frequency of NAcSh-projecting IL neurons in SH mice is comparable to that of GH neurons (Two-tailed unpaired t test with Welch's correction, $t_{28.36}$ = 0.2572, $P$ = 0.7989, n = 14 cells from 2 GH mice, n = 18 cells from 4 SH mice).

**d,** Input resistance of NAcSh-projecting IL neurons in SH mice is comparable to that of GH mice (Two-tailed unpaired t-test with Welch's correction, $t_{67}$= 1.63, $P$ = 0.1079 n = 32 cells from 14 GH mice, n = 46 cells from 13 SH mice,).

**e,** Resting membrane potential (RMP) of NAcSh-projecting IL neurons in SH mice is comparable to that of GH mice (Two-tailed unpaired t-test with Welch's correction, $t_{49.35}$= 1.281, $P$ = 0.2060, n = 15 cells from 14 GH mice, n = 29 cells from 13 SH mice).

**f,** Membrane capacitance ($C_m$) of NAcSh-projecting IL neurons in SH mice is comparable to that of GH mice (Two-tailed unpaired t-test with Welch's correction, $t_{84.95}$= 1.538, $P$ = 0.1279, n = 41 cells from 14 GH mice, n= 52 cells from 13 SH mice).

**g,** Action potential threshold of NAcSh-projecting IL neurons in SH mice is comparable to that of GH mice (Two-tailed unpaired t-test with Welch's correction, $t_{81.67}$= 0.9535, $P$ = 0.3431, n = 40 cells from 14 GH mice, n = 51 cells from 13 SH, ).

**h,** Rheobase current of NAcSh-projecting IL neurons in SH mice is comparable to that of GH mice (Two-tailed unpaired t-test with Welch's correction, $t_{85.53}$= 1.054, $P$ = 0.2950, n = 40 cells from 14 GH mice, n = 52 cells from 13 SH mice, ).

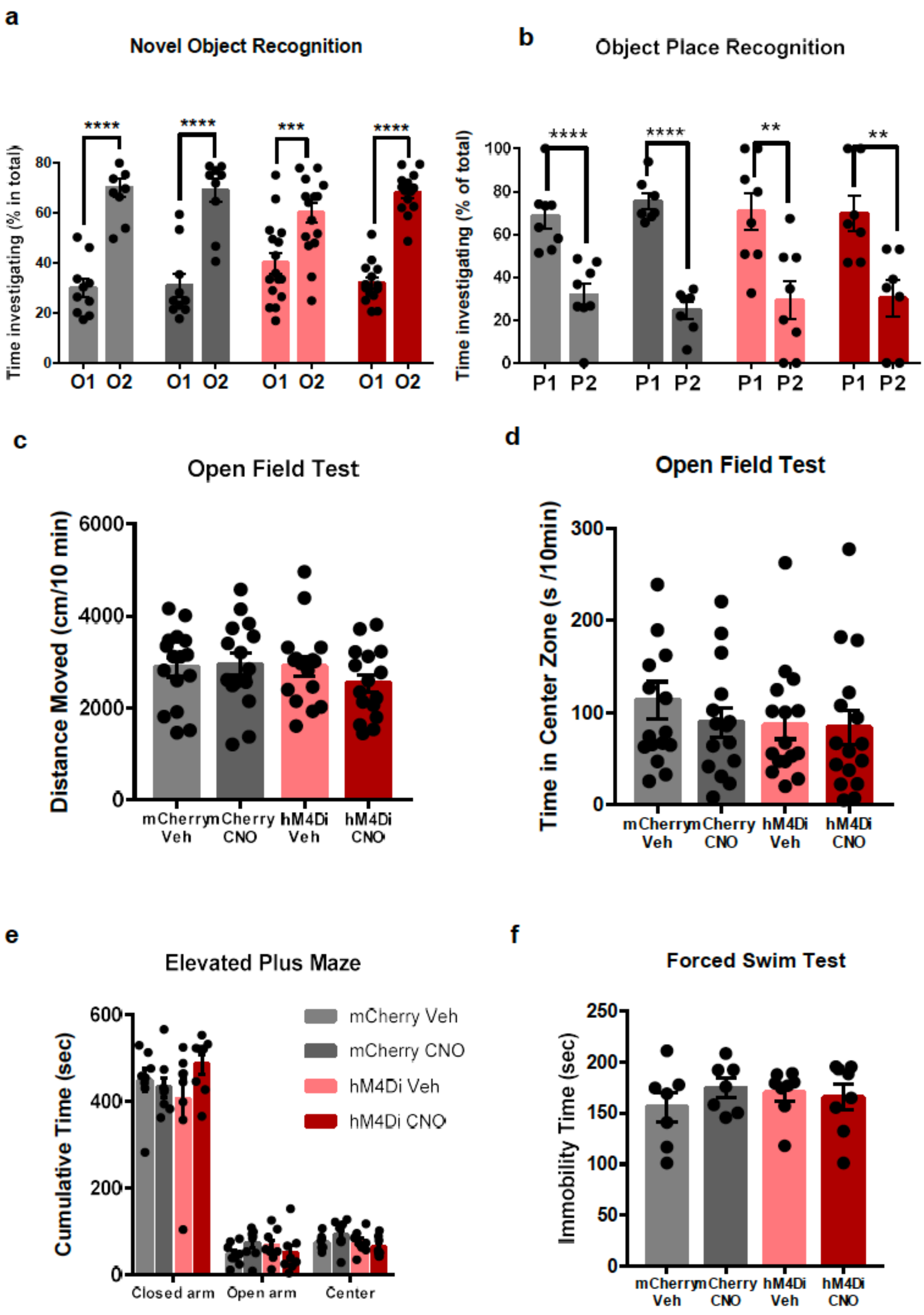


**Extend Data figure 5. Chemogenetic inhibition of NAcSh-projecting IL neurons did not affect either basal locomotor activity or anxiety level in GH mice.**

**a,** Inactivating NAcSh-projecting IL neurons does not hamper the novel object recognition (n = 16 mice (mCherry Veh), 15 mice (mCherry CNO), 16 mice (hM4Di Veh), 15 mice (hM4Di CNO), Two-tailed paired t-test: ****P* = 0.0003 (mCherry Veh), ***P* = 0.0022 (mCherry CNO), **P* = 0.0264 (hM4Di Veh),

****$P$ < 0.0001 (hM4Di CNO). Veh, vehicle.

**b,** Inactivating NAcSh-projecting IL neurons does not impair hippocampal-dependent memory in object place recognition test. (n = 8 mice (mCherry Veh), 7 mice (mCherry CNO), 8 mice(hM4Di Veh), 7 mice(hM4Di CNO), two-way ANOVA $F_{1, 26}$ = 2.117, $P$ = 0.1576, [between-group comparison by Sidak's multiple comparisons test: mCherry Veh, $P_1 - P_2$ $t_{36}$ = 5.444, ****$P$ < 0.0001, mCherry CNO, $P_1 - P_2$ $t_{36}$ = 7.085, ****$P$ < 0.0001], two-way ANOVA $F_{1, 26}$ = 0.008692, $P$ = 0.9264, [between-group comparison by Sidak's multiple comparisons test: hM4Di Veh, $P_1 - P_2$ $t_{26}$ = 3.488, **$P$ = 0.0035, hM4Di CNO, $P_1 - P_2$ $t_{26}$ = 3.135, **$P$ = 0.0084]).

**c,** Inactivating NAcSh-projecting IL neurons does not affect the locomotive activity of mice (n = 16 Control Veh, n = 15 Control CNO, n = 16 hM4Di Veh, n = 16 hM4Di CNO, ordinary one-way ANOVA, F = 0.788 , $P$ = 0.5054).

**d**, Inactivating NAcSh-projecting IL neurons does not affect the time spent at the center zone in the open field test (n = 16 Control Veh, n = 15 Control CNO, n = 16 hM4Di Veh, n = 16 hM4Di CNO, ordinary one-way ANOVA, F = 0.6135 , $P$ = 0.6089).

**e,** Inactivating NAcSh-projecting IL neurons does not affect the anxiety level of mice tested in elevated plus maze (n = 8 Control Veh, n = 8 Control CNO, n = 8 hM4Di Veh, n = 8 hM4Di CNO, ordinary one-way ANOVA, F = 0.004346 , $P$ = 0.9996).

**f,** Inactivating NAcSh-projecting IL neurons does not affect the immobility time in the forced swim test (n = 7 Control Veh, n = 7 Control CNO, n = 8 hM4Di Veh, n = 8 hM4Di CNO, ordinary one-way ANOVA, F = 0.4852 , $P$ = 0.6955).

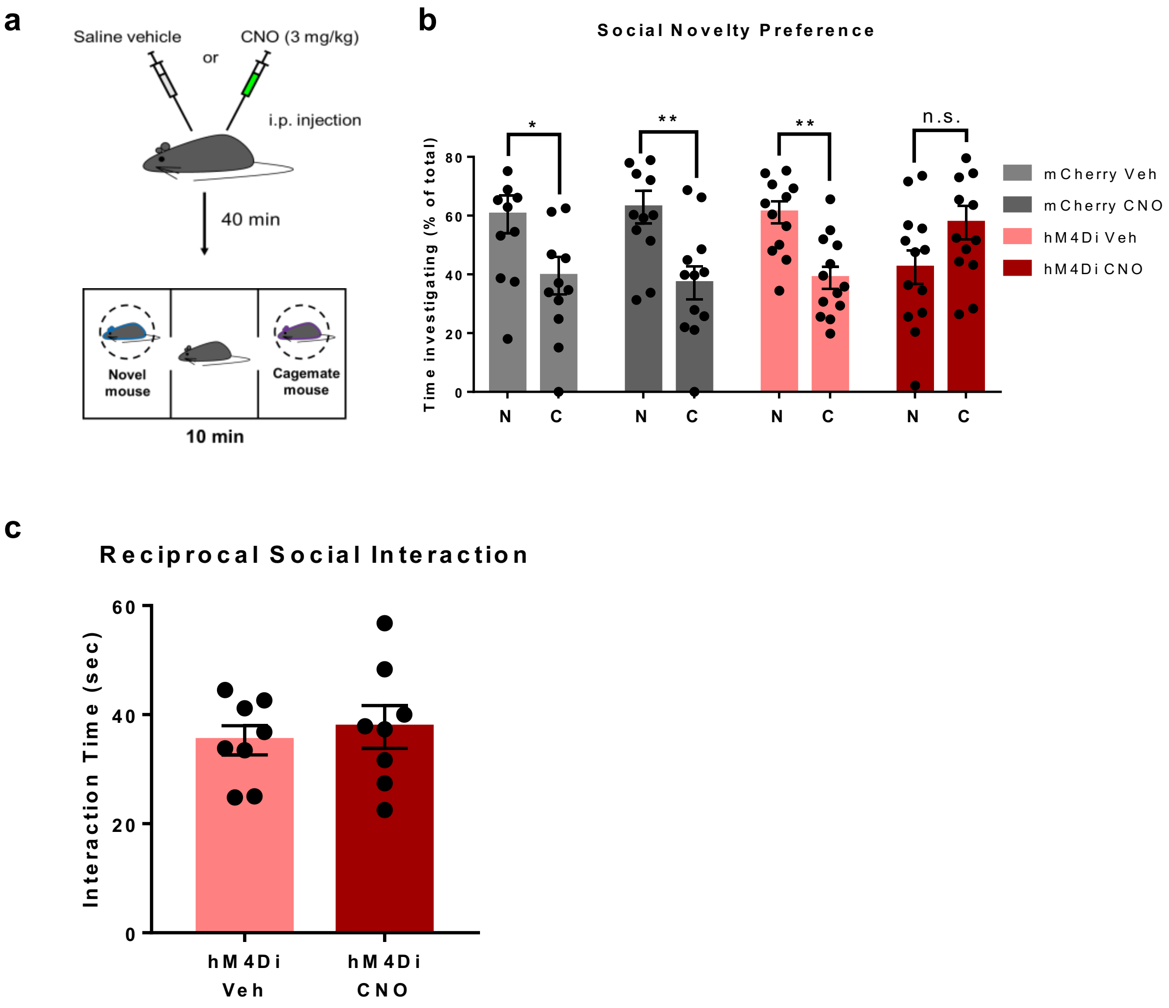


**Extend Data figure 6. Assessments of sociability in chemogenetic inhibition of NAcSh-projecting IL neurons by using cage mate as a target mouse.**

**a,** Schematic diagram for social novelty preference test with a novel mouse and a cage mate as a target mouse.

**b**, Inhibition of NAcSh-projecting IL neurons impairs social recognition even when a cage mate is used as a target mouse. (n = 12 mice (mCherry Veh), 12 mice (mCherry CNO), 13 mice(hM4Di Veh), 13 mice(hM4Di CNO), two-way ANOVA $F_{1, 44}$ = 0.1669, $P$ = 0.6849, [between-group comparison by Sidak's multiple comparisons test: mCherry Veh, N – C $t_{44}$ = 2.453, *$P$ = 0.0361, mCherry CNO, N – C $t_{44}$ = 3.031, **$P$ = 0.0081], two-way ANOVA $F_{1, 48}$ = 15.05, ***$P$ = 0.0003, [between-group comparison by Sidak's multiple comparisons test: hM4Di Veh, N – C $t_{48}$ = 3.259, **$P$ = 0.0041, hM4Di CNO, N – C $t_{44}$ = 2.228, $P$ = 0.0603])

n.s., not significant.

**c**, Chemogenetic inhibition of NAcSh-projecting IL neurons did not impair social interaction examined in reciprocal social interaction test. Two-tailed unpaired t-test with Welch's correction, $t_{12.34} = 0.5187$, $P = 0.6132$, n = 16 biologically independent mice interacted as 8 pairs.
